# Cis-TERN: a graph-based method to integrate single-cell chromatin accessibility with gene expression data to infer the pseudotime ordering of cells

**DOI:** 10.1101/2025.11.21.689849

**Authors:** Ameya Gokhale, Pawel F. Przytycki

## Abstract

Pseudotime ordering methods computationally arrange individually sequenced cells along a continuous trajectory to infer their progression through biological processes. Chromatin accessibility at regulatory elements is pivotal for orchestrating the timing of gene expression during cellular differentiation. However, the sparsity inherent in single-cell data, along with the discrete grouping imposed by clustering-based approaches, often obscures subtle cell-state transitions and intermediate populations. Here, we introduce Cis-TERN (Cis-Temporally Encoded Regulation Network), a graph-based method that integrates single-cell measurements of gene expression (scRNA-seq) and chromatin accessibility (scATAC-seq) to infer a pseudotime ordering of cells. Exploiting the fact that distal *cis* chromatin accessibility often precedes gene activation, Cis-TERN builds a directed bipartite graph between scRNA-seq and scATAC-seq cells with edge weights determined by either proximal or distal accessibility. Through network diffusion, Cis-TERN assigns continuous pseudotime scores to all cells. Cis-TERN infers cellular pseudotime without the need for prior biological knowledge, such as specifying a root cell type, thereby enabling unbiased reconstruction of differentiation from data alone. We show that our scores align with known lineages in healthy and tumor-derived hematopoietic cells. Additionally, Cis-TERN is able to trace individual cancer cells back to progenitor cell types. Our approach establishes a framework for leveraging chromatin accessibility and gene expression jointly to learn the timing of cell state progression.

## 1. Introduction

Chromatin accessibility at regulatory elements plays a major role in dynamic control of gene expression determining both its timing and specificity [1], [2]. By selectively allowing for transcription factors to bind to enhancer and promoter regions, cells launch the transcriptional programs suited to their development [1], [3]. Changes in accessibility drive cellular differentiation [2], [3], [4]. In diseases such as cancer, tumors often exhibit abnormal chromatin accessibility patterns resulting in irregular differentiation [4], [5]. Understanding chromatin accessibility and its relation to gene expression is therefore essential for determining the mechanism behind the progression of normal healthy cells into diseased ones. While chromatin accessibility proximal to and overlapping genes is generally required for them to be expressed and is therefore thought to be concurrent with a gene being expressed, distal accessibility at regulatory elements indirectly modulates the magnitude of expression [1], [2], [3], [6], [7]. Crucially, distal peak accessibility often precedes gene expression and therefore can be a determinant of lineage selection [8].

Advances in high-throughput sequencing technology have allowed for the measurement of chromatin accessibility (scATAC-seq) and gene expression (scRNA-seq) profiles in individual cells [2], [6]. The resulting data has allowed researchers to better understand cell differentiation in a computationally tractable manner called pseudotime ordering where individual cells are ordered across differentiation [9], [10]. However, because datasets generated from these techniques, especially scATAC-seq, suffer from sparsity issues, cells are typically aggregated into discrete clusters for analysis, which are interpreted as cell types or states [10]. Pseudotime inference typically models cellular differentiation as a sequence of transitions between these clusters or alternative lower dimensionality aggregations such as trajectories [10]. For example, Monocle 3, a commonly used graph-based method for pseudotime inference from scRNA-seq data, first builds a clustered graph and then fits a principal curve through those cluster partitions, assigning pseudotime first as the geodesic distance from a user-chosen root cell or cluster without further explicit consideration of each individual cell [9], [11]. Some pseudotime methods incorporate chromatin accessibility, but either treat it in isolation rather than integrating it with transcriptomics or restrict attention to proximal sites to convert accessibility into gene scores for direct integration with scRNA-seq data [12], [13]. These methods do not explicitly take advantage of the orthogonal information provided by chromatin accessibility. Additionally, most pseudotime inference methods do not infer directionality and require manual specification of the cell type of origin [10].

To address these shortcomings, we introduce Cis-TERN (Cis-Termporally Encoded Regulation Network), a graph-based method to infer timing of differentiation in scRNA-seq data through integration with scATAC-seq data. Cis-TERN is designed to exploit the observation that distal *cis* chromatin is often accessible before a target gene is expressed. To do this, Cis-TERN constructs a bipartite graph between scRNA-seq and scATAC-seq cells with directed edges encoding either proximal or distal accessibility-based similarity between cells. Through network diffusion, it then assigns a continuous pseudotime score to all cells. We validated Cis-TERN on scRNA-seq and scATAC-seq datasets from healthy hematopoietic cells, demonstrating that Cis-TERN orders cells in a manner that is consistent with known biology and additionally found that Cis-TERN is able to order cancer cells per-patient in mixed-phenotype acute leukemia (MPAL) samples [14]. Finally, the Cis-TERN graph can be used to trace individual cells back to progenitor cell types. We applied this “reverse tracing” to terminal blood cell types to trace them back to hematopoietic stem cells (HSCs) and to cancer cells in the MPAL data to trace their progression from healthy cells.

## 2. Results

### 2.1. Overview of method

Cis-TERN is a graph-based method that generates per-cell pseudotime scores by constructing a bipartite graph whose nodes are scATAC-seq and scRNA-seq cells. Nodes are connected by weighted, directed edges. Edges from scRNA-seq cells to scATAC-seq cells encode cross modal similarity, computed as the cosine similarity between the scRNA-seq cell’s gene expression profile and the scATAC-seq cell’s proximal and gene-body accessibility (Figure 1, grey edges). Edges from scATAC-seq cells to scRNA-seq cells are weighted by a precedence score (Figure 1, green edges), derived from the correlation between distal *cis* peak accessibility and gene expression, reflecting the expectation that accessibility at cis regulatory elements often precedes transcription. For each such edge, the precedence score combines (i) the scATAC-seq cell’s chromatin accessibility, (ii) the scRNA-seq cell’s gene expression levels, and (iii) inferred peak-gene regulatory links (see Methods section 4.1 for details). Given the sparsity and noisiness of single-cell data, individual edges will carry limited timing information. To aggregate the timing signal globally and smooth over noise, Cis-TERN performs a random walk with restart on the bipartite graph and uses the stationary probability of each scRNA-seq node as its pseudotime score. Intuitively, even weak precedence signals accumulate over long walks, biasing the stationary distribution toward later (higher-pseudotime) cells (see Methods section 4.2 for details).

**Figure 1.**
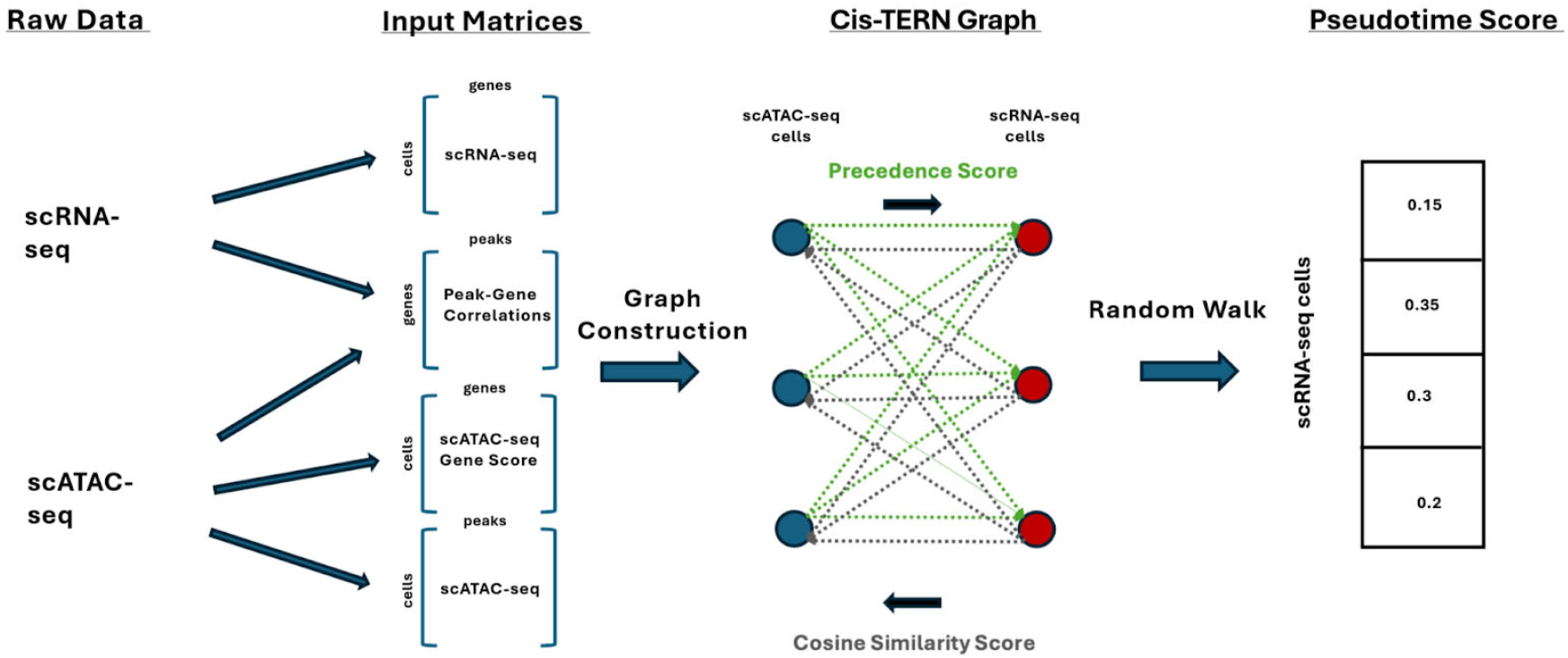
Overview of Cis-TERN. Cis-TERN takes as input four matrices generated from scRNA-seq and scATAC-seq data to construct a directed bipartite graph with edges weighted by cosine similarity in one direction and precedence score in the other on which it performs a random walk to calculate pseudotime scores for each scRNA-seq cell.

Cis-TERN also enables “reverse tracing” individual cells to their progenitors. To do so, Cis-TERN reverses the precedence edge weights in the graph and again runs a random walk to produce a reverse pseudotime score per scRNA-seq cell, where higher values indicate cells more likely to appear earlier in pseudotime. Cis-TERN then constructs a second graph over scRNA-seq cells alone, with directed edges weighted by cosine similarity of gene-expression profiles. It then retains, for each cell, only its highest scoring connections stratified by reverse pseudotime (Appendix Figure 1). A Monte Carlo path sampling procedure outputs the most probable differentiation trajectory for any chosen scRNA-seq cell (see Methods section 4.3 for details).

### 2.2. Cis-TERN recapitulates known lineages in hematopoiesis

We tested Cis-TERN’s functionality on scRNA-seq (35,882 cells) and scATAC-seq (36,690 cells) data from healthy bone marrow and peripheral-blood mononuclear cells (PBMCs) and six samples of mixed phenotype acute leukemia (MPAL) consisting of a total of 18,056 scRNA-seq cells and 35,423 scATAC-seq cells also collected from bone marrow and peripheral blood [14].

To assess Cis-TERN’s ability to assign pseudotime scores to individual cells, we first tested whether it recapitulates established differentiation lineages in the scRNA-seq and scATAC-seq cells from the healthy individuals. The scRNA-seq had previously been clustered into 26 hematopoietic states including lineages for monocytes and B cells. We ran Cis-TERN on the full combined scRNA-seq and scATAC-seq data to infer pseudotime scores for all cells in both lineages [2], [3]. As expected, Cis-TERN scores increase monotonically on average across successive cell types in both known lineages (Figure 2a for monocytes and Figure 2b for B cells) with the exception of HSCs, which have an inflated number of incoming non-zero edges increasing their precedence score (Appendix Figure 2a). For scores for all cell types, see Appendix Figure 3. We additionally found that Cis-TERN is able to order cell types in the developing brain, an organ with complex lineages [15], [16] (Appendix Figure 4). Overall, this indicates that Cis-TERN is able to recapitulate the broad global ordering of cell lineages with no prior knowledge.

**Figure 2.**
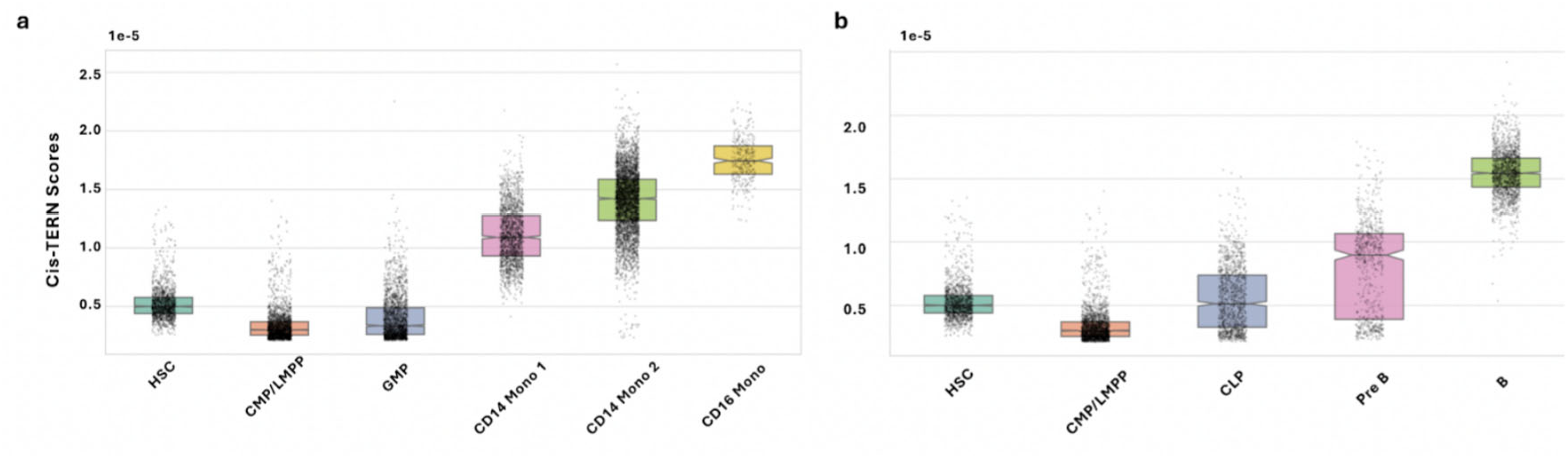
Cis-TERN orders cells from known lineages in hematopoiesis. Notched boxplots of the Cis-TERN pseudotime scores for cells annotated to cell types ordered by their lineages for **a)** monocytes and **b)** B cells.

**Figure 3.**
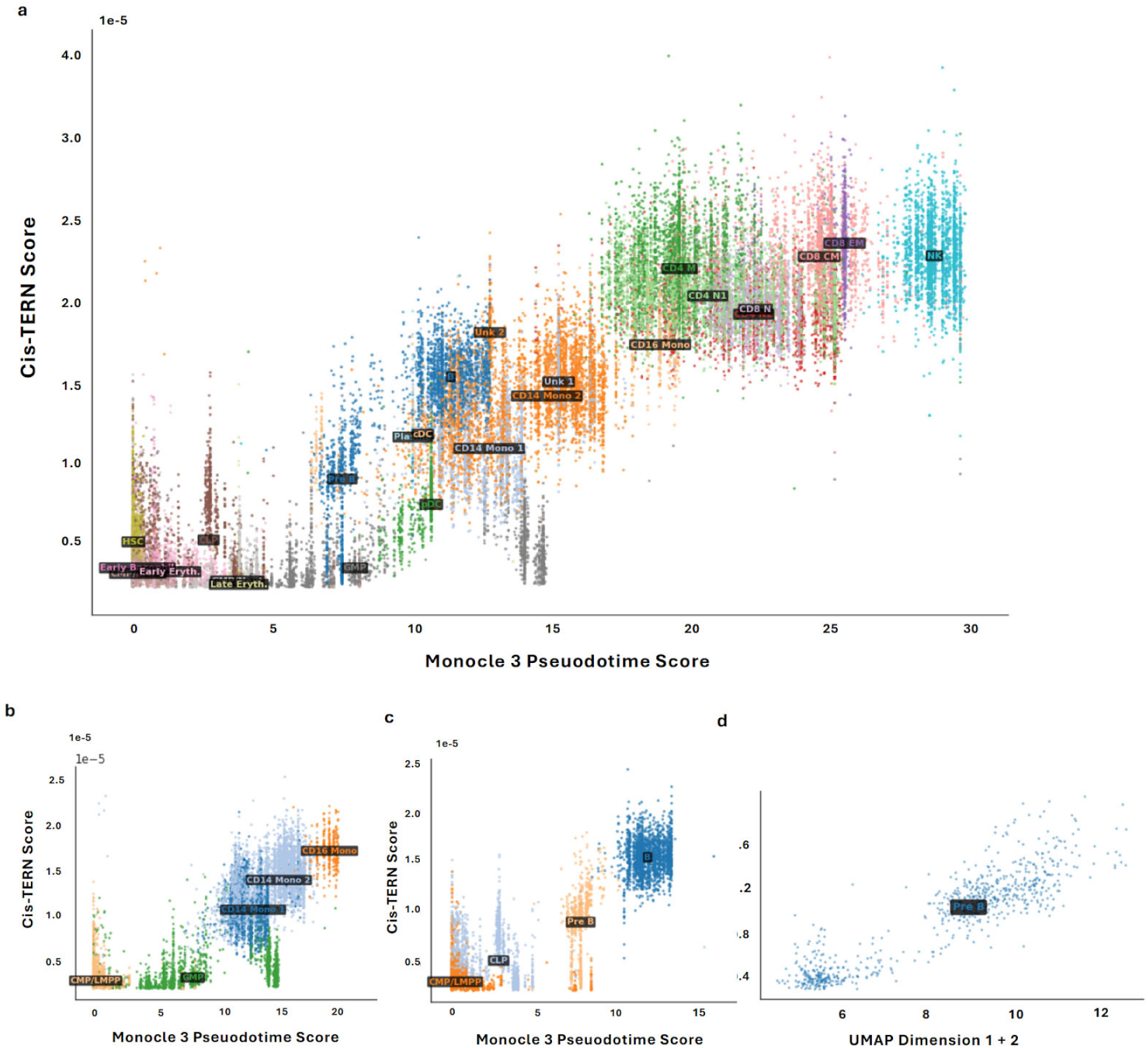
Cis-TERN captures the continuous progression of cells. Cis-TERN scores are highly correlated to Monocle 3 pseudotime scores across **a)** all scRNA-seq cells **b)** the monocyte Lineage and **c)** the B-cell lineage. **d)** Cis-TERN scores for pre-B cells are highly correlated with the UMAP embedding of scRNA-seq PBMC data [14]. In all panels, cells are colored based on annotated cell type labels and cell type labels appear at the centroid of all cells assigned to that cell type.

**Figure 4.**
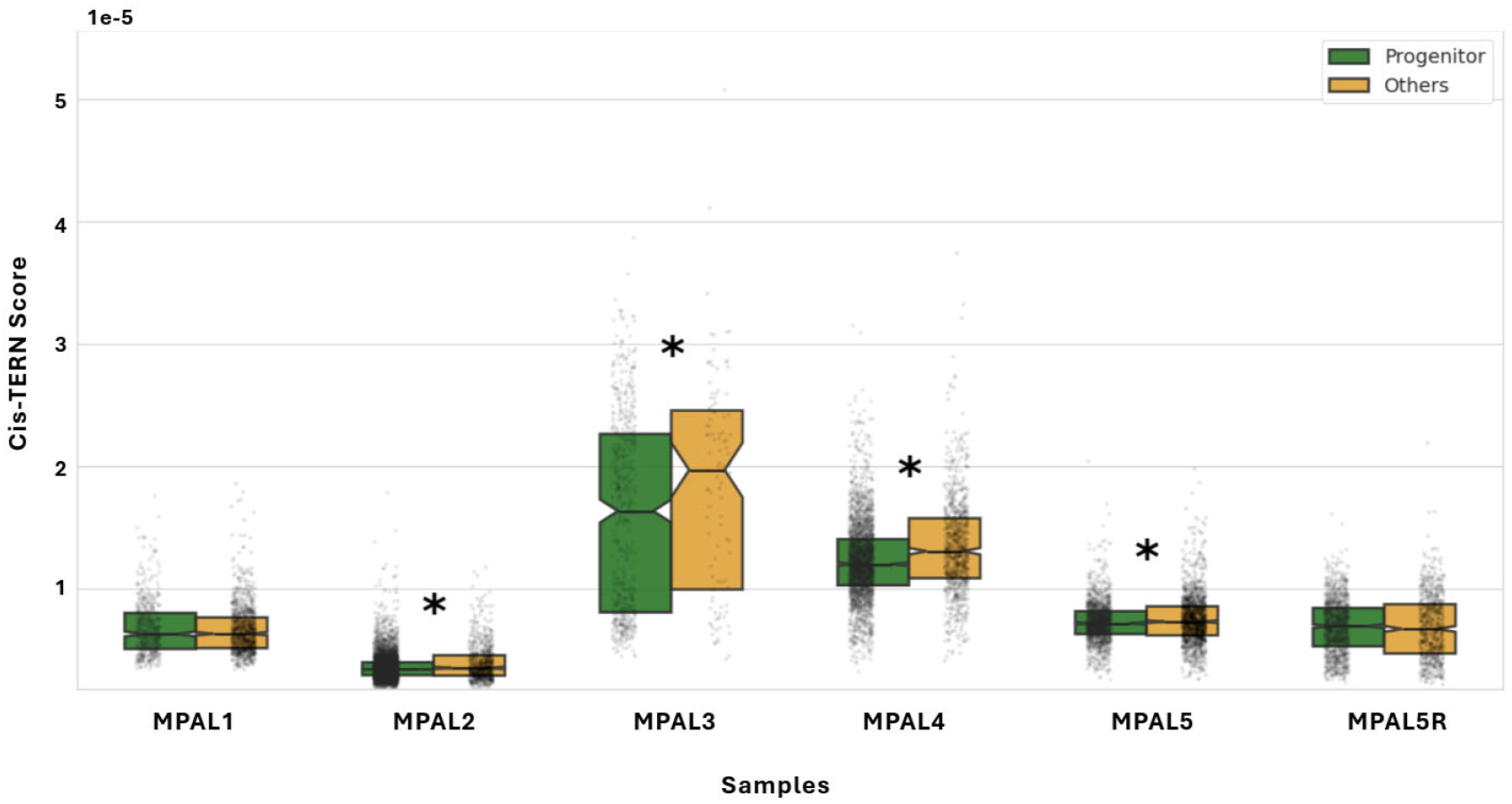
Cis-TERN differentiates progenitor-like cancer cells. For four of six tumor samples, progenitor-like cells (green notched boxplots) had significantly lower Cis-TERN scores than other cancer cells (orange notched boxplots). Significance level of a one-tailed Wilcoxon test of *p*-value <0.05 is indicated with a *.

Next, we examined how Cis-TERN captures the continuous progression of cells both across cell types and within cell types. Since a ground truth is not known, we compared Cis-TERN’s pseudotime scores to those of Monocle 3 [9] and to the ordering captured by a low dimensional UMAP embedding can capture local cell type similarity and progression, including in hematopoiesis [14], [17]. When running Monocle 3 on integrated scRNA-seq and scATAC-seq data with HSCs set as the root, we observed that, across all cells, Cis-TERN’s pseudotime scores correlate well those generated by Monocle 3 despite Cis-TERN having no prior knowledge of a cell type of origin (Figure 3a, Pearson’s p=0.89). We similarly observed that within the monocyte and B-cell lineages, the two models broadly agreed (Figure 3b the for monocyte lineage, Pearson’s p=0.77 and Figure 3c for the B-cell lineage, Pearson’s p=0.85). In both lineages, Cis-TERN showed a better ability to assign a broad range of scores to cells within the same annotated cell type and generated smoother, less discretized transitions between cell types. To further evaluate the local ordering of our scores, we compared them with a UMAP embedding of the scRNA-seq data (Appendix Figure 5a) [14], [17]. We observed that Cis-TERN’s pseudotime scores correlate well with the embedding across both lineages (Appendix Figure 5b, Pearson’s p=0.60 for the monocyte lineage and Appendix Figure 5c, Pearson’s p=0.81 for the B-cell lineage). We similarly found that, within each cell type, our scores correlated with the embedding, including for transitionary cell types such as pre-B cells (Figure 3d, Pearson’s p=0.90) and for cell types that appear more continuous than discrete such as CD14 monocytes types 1 and 2 (Appendix Figure 5d, Pearson’s p=0.49). Additionally, while, as observed above, the median HSC Cis-TERN score was not the lowest among all cell types, the local gradient is in agreement with the embedding highlighting our method’s ability to resolve progressive differentiation within the population (Appendix Figure 5e, Pearson’s p=0.27).

**Figure 5.**
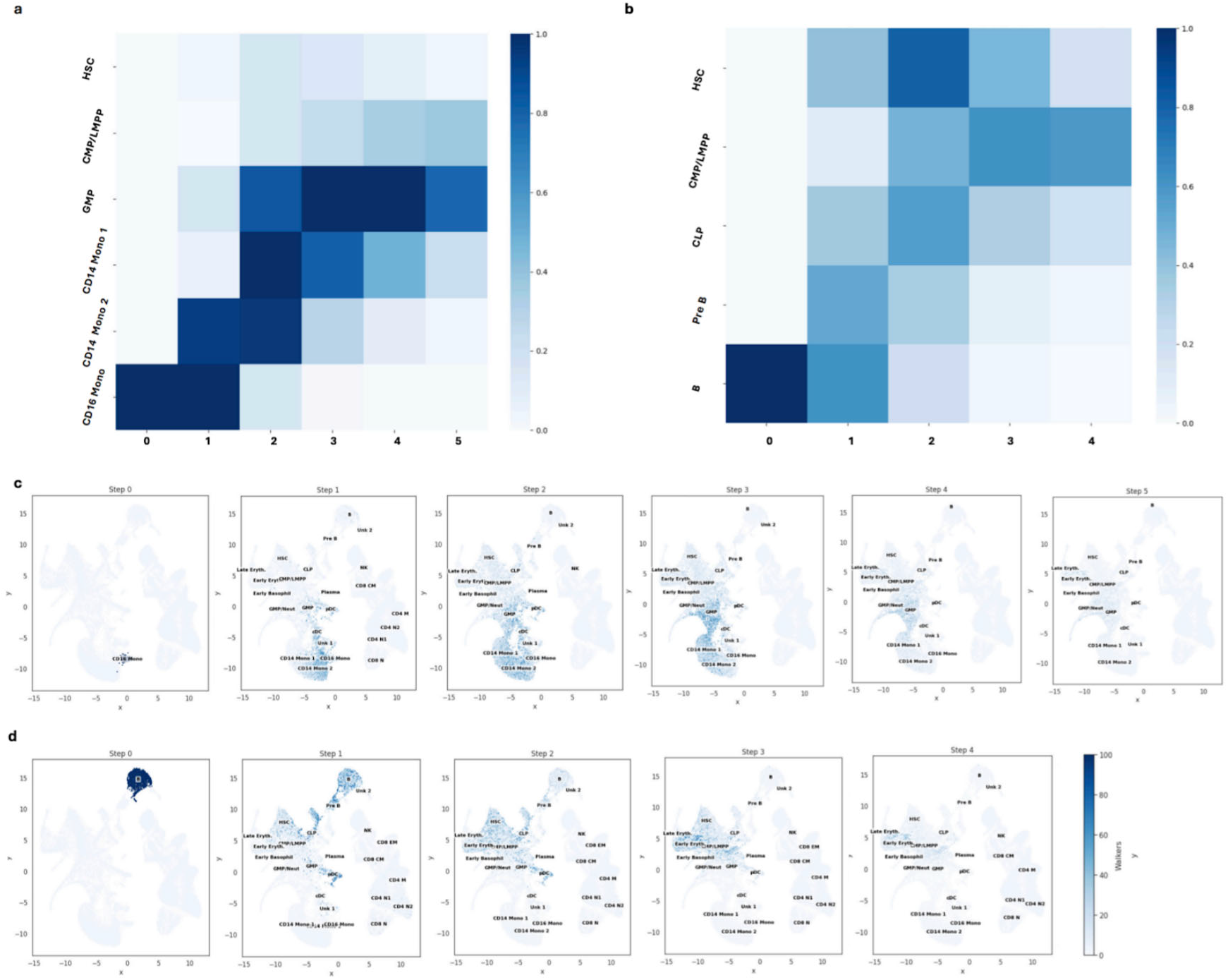
Reverse tracing terminal cell types to progenitors. The min-max scaled fraction of traces that visited each cell type in **a)** the monocyte and **b)** the B-cell lineage when all traces starting at the corresponding terminal cell type are aggregated. The number of individual traces starting at the terminal cells for **c)** the monocyte and **d)** the B-cell lineage observed to be at every cell in the UMAP embedding of the scRNA-seq data at each time step [14].

Given that scATAC-seq data is typically less readily available than scRNA-seq data [18], we next examined robustness of Cis-TERN to smaller quantities of scATAC-seq cells. We ran Cis-TERN on the full PBMC scRNA-seq dataset described above while holding out all but one scATAC-seq sample at a time. This captures the common experimental practice of performing scRNA-seq on all samples but scATAC-seq on only a subset [18]. In this setting, Cis-TERN recapitulated lineage progression in both B cells and monocytes trajectories with as around 10,000 scATAC-seq cells. At substantially lower coverage (i.e., fewer than 5,000 scATAC-seq cells), performance degraded, suggesting a practical lower bound on input size for reliable reconstruction (Appendix Figure 6). Across these samples, Cis-TERN completed scoring in around 45 seconds for samples with around 3,000 cells to just over 2 minutes for the full dataset with over 35,000 cells per modality, indicating favorable runtime scaling (Appendix Figure 7). As an additional robustness test, we examined the performance of Cis-TERN using precedence scores or cosine similarities alone. To evaluate precedence scores, we took their sum per scRNA-seq cell and observed that they were able to partially order the lineages alone (Appendix Figure 8a and 8b) and were highly correlated with Cis-TERN scores per cell type (Appendix Figure 8c). To evaluate cosine similarity alone, we replaced precedence weights with cosine similarity in the bipartite graph which resulted in complete loss of ordering of cell types in lineages as expected (Appendix Figure 8d and 8e).

**Figure 6.**
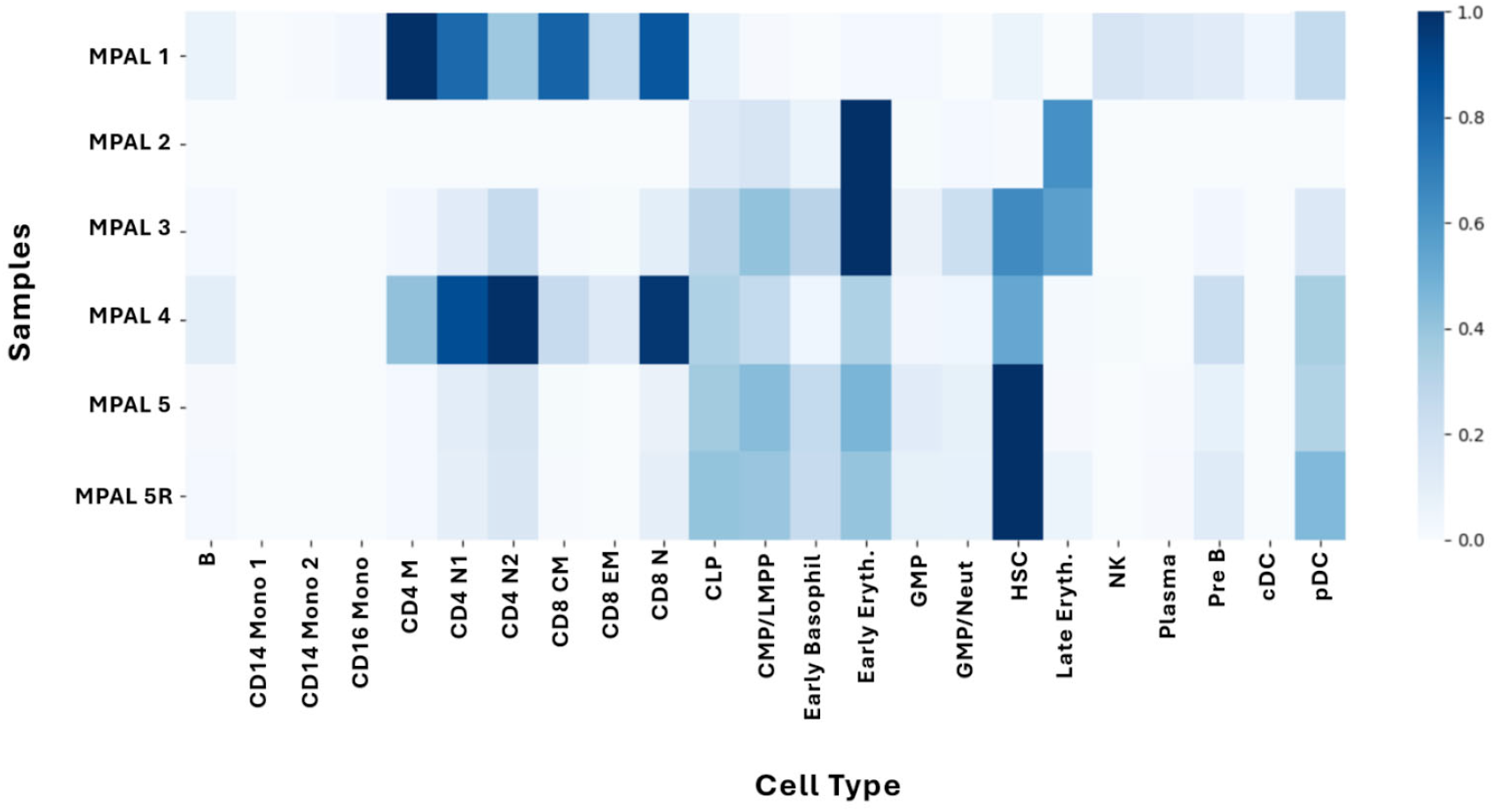
Reverse tracing cancer cells reveals tumor heterogeneity. The fraction of trace that ends at each cell type after three steps when starting at cancer cells in the given sample.

Finally, we used Cis-TERN to pseudotime order cancer cells from MPAL tumor samples in a per patient manner. The six MPAL samples were classified as T-myeloid MPAL (samples “MPAL1” - “MPAL3”), B-myeloid MPAL (sample “MPAL4”), T-myeloid MPAL sampled before CALGB chemotherapy (sample “MPAL5”), and post-treatment relapse (sample “MPAL5R”) [14]. To score each sample, we ran Cis-TERN on the combined full healthy hematopoiesis data with the scRNA-seq and scATAC-seq data for that sample (see Methods section 4.4 for details). First, we observed that the ordering of the healthy cells was largely unchanged by adding the tumor data as would be expected (Appendix Figure 9). Across the tumor samples, while there is no ground truth for the time ordering of cancer cells, we observed that cells that had been annotated as “progenitor like” had earlier Cis-TERN pseudotime scores (Figure 4). This indicates that Cis-TERN was broadly able to, in individual samples, sort cancer cells by their plasticity.

### 2.3. Cis-TERN reverse tracing retrieves progenitor paths for terminal state cells

In order to evaluate Cis-TERN’s capacity to “reverse trace” individual cells to their progenitors, we ran Cis-TERN’s reverse tracing procedure on the terminal cell types in healthy hematopoiesis. In particular we ran it on all 256 CD16 monocytes (the terminal cell type of the monocyte lineage) and on all 1,711 B cells (the terminal cell type of the B-cell lineage). For each terminal type, we then aggregated the traces from all the cells and observed that the majority of traces moved back through the lineage to progenitor cell types (Figure 5a for CD16 monocytes and Figure 5b for B cells) though the traces do make additional spurious visits to other cell types as would be expected to occur due to stochastic variability (Appendix Figure 10). The number of steps taken before termination is automatically determined as described in Methods section 4.3 (See Appendix Figure 11 for calculated distribution of walk lengths). To explore global per-cell reverse tracing dynamics, we used the precomputed UMAP of scRNA-seq cells to observe the cells visited by the traces in both lineages

(Figure 5c for CD16 monocytes and Figure 5d for B cells). As expected, the density of the traces moves gradually towards progenitor cell types.

Finally, given Cis-TERN’s ability to retrace the paths of healthy terminal state cells, we tested how Cis-TERN would reverse trace cancer cells to possible healthy progenitors. We applied the reverse tracing procedure to the per-sample integrations described in section 2.2 aggregating the traces from all cancer cells in each sample and observed that cancer cells had sample specific origins (Figure 6) to which traces converge to after a small number of steps (Appendix Figure 12). Cancer cells from two samples (MPAL1 and MPAL4) traced primarily back to T cells, while two samples (MPAL2 and MPAL3) had similarity to erythroid cells, as was previously observed [14]. Finally, MPAL5 and MPAL5R, which were taken from the same individual prior to chemotherapy and after treatment and relapse, showed very similar cancer origins as would be expected and was previously observed [14].

## 3. Discussion

Cis-TERN provides a framework for integrating scRNA-seq and scATAC-seq measurements to infer cellular differentiation by leveraging the precedence of distal chromatin accessibility. Our approach enables the reconstruction of developmental progressions without prior knowledge of root cell states and successfully aligns with established biological lineages in healthy hematopoiesis. Additionally, Cis-TERN retains favorable scaling to tens of thousands of cells per modality. Beyond forward ordering, Cis-TERN’s unique reverse tracing allows individual cells to be mapped back to their putative progenitors. We demonstrate this in the context of mixed-phenotype acute leukemia where Cis-TERN mapped malignant cells to which healthy precursors they likely differentiated from.

While Cis-TERN was largely able to order developmental lineages, it consistently placed HSCs later than expected. Though we found that this was due to an inflated number of non-zero incoming precedence edges to these cells, it is unclear why these edges occurred. One possible explanation is that HSCs may have more within-cell type precedence edges due to being self-renewing stem cells.

A core component of Cis-TERN are the peak-gene link predictions used to find distal *cis* regulatory elements to associate with each gene for calculating precedence scores. The current implementation of Cis-TERN uses across-cell-type correlation between scRNA-seq and scATAC-sec data as an indication of linkage. However, this is a coarse-grained view that only captures the subset of links that is present in common cell types. In the future, the integration of additional data modalities such as H3K27ac and Hi-C could enhance link prediction to generate better precedence scores by identifying active enhancers and enhancer-promoter contacts.

Overall, Cis-TERN is an immensely flexible framework for integrating scATAC-seq and scRNA-seq data. The encoding of this multimodal data into a weighted bipartite graph leaves room for alternative edge weights and scales well to large quantities of single-cell sequencing data.

## 4. Methods

Cis-TERN (Cis-Termporally Encoded Regulation Network) generates a pseudotime ordering of cells by integrating scRNA-seq and scATAC-seq data by building a graph (described in section 4.1) and then walking on the graph to score cells (described in section 4.2). It can trace cells back to their origins (described in section 4.3). Datasets used, data processing, and details of how the method was run are described in section 4.4. Code availability is described in section 4.5.

### 4.1 Generating Cis-TERN graph

As input, Cis-TERN takes a cell-by-gene expression matrix *S*^*G*^, a cell-by-peak accessibility matrix *S*^*A*^, a cell-by-gene gene-score matrix *S*^*M*^ and a gene-by-peak peak-gene correlation matrix *S*^*L*^ [7]. These matrices can be generated from one or more scATAC-seq and scRNA-seq experiments following the data processing steps described in section 4.4. Note that the cells in *S*^*G*^ do not need to be the same as the cells in *S*^*A*^. Using these matrices, Cis-TERN generates a directed bipartite graph where nodes are scATAC-seq and scRNA-seq cells. There are two types of directed edges between each group of cells as follows:

#### Precedence Score Edges

These are directed edges from scATAC-seq to scRNA-seq cells. A precedence score between an scATAC-seq cell and scRNA-seq quantifies how strongly the scATAC-cell precedes the scRNA-seq cell based on distal accessibility of peaks *cis* to expressed genes. We calculate the cell-by-cell matrix of precedence scores *P* in the following manner:

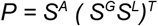

Some values in *P* will be negative due to negative correlations between peaks and genes. These values are set to 0.

#### Cosine Similarity Score Edges

These are directed edges from scRNA-seq to scATAC-seq cells. A cosine similarity score captures how similar each cell’s gene expression in the scRNA-seq data is to proximal and gene body accessibility of cells in the scATAC-seq data using cosine similarity (for details of how accessibility was calculated per gene, see section 4.4). This results in a cell-by-cell matrix *C* where each entry *C*_*i*,,*j*_ is the cosine similarity of cell *i* in the scATAC-seq data and cell j in the scRNA-seq data across genes.

The final Cis-TERN graph can be written as *G = (V, E)* where *V* is all scATAC-seq cells and scRNA-seq cells and *E* is the two sets of directed precedence and cosine similarity edges. We represent this graph with the square adjacency matrix *A* where the number of rows and columns equal to the total number of cells and where the top right quadrant is *P* and bottom left quadrant is *C*^*T*^.

### 4.2 Cis-TERN pseudotime scoring

To capture temporal progression, Cis-TERN performs a random walk with restart. We first make the adjacency matrix *A* column stochastic. To allow for restarts, we then calculate the transition matrix *G* as follows:

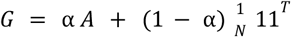

where *N* is the total number of nodes in the graph (scATAC-seq + scRNA-seq cells), 11^𝑇^ is a matrix of all ones (used to uniformly transport a node for a restart), and α is the damping factor that controls the probability of teleportation. Adding a restart probability guarantees the existence of a unique steady state distribution which is a vector of probabilities π satisfies π = *G*π.

Cis-TERN uses an iterative method to find the steady state distribution by initializing π_0_ to be uniform and then iterating:

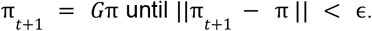

Upon convergence, the final vector π^*^ is an approximation of the true underlying steady state distribution. Each scRNA-seq cells value in π^*^ is its Cis-TERN score. Intuitively, a higher score means a random walker will visit that cell more in the long run, implying a later position in pseudotime.

### 4.3 Cis-TERN reverse tracing

To trace backward from a given scRNA-seq cell and infer its putative progenitors, we modify the Cis-TERN bipartite graph by inverting the precedence weights giving a reverse precedence score 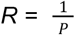 In the original graph, a larger precedence weight on edge A to B implies that B lies later in pseudotime than A. After inversion, this temporal signal is reversed: higher weights now favor transitions to cells that are earlier in development.

Cis-TERN then constructs an adjacency matrix *A\** where the top right quadrant is *R* and bottom left quadrant is *C*^*T*^. We run the same Cis-TERN random walk algorithm described in 4.2 with this new graph. In this case a higher random walk score indicates that the cell came earlier in development trajectory with the final steady state vector *t* capturing a “reverse Cis-TERN score” for the scRNA-seq cells. Cis-TERN next builds a pairwise cosine similarity matrix *S*^*S*^ from *S*^*G*^ only keeping the top 10% of entries by cosine similarity in each row (scRNA-seq cell) and setting the rest to 0. Additionally, for each row, Cis-TERN only keeps the edges to other cells that have a greater reverse Cis-TERN score .

Finally, to trace individual cells back to their progenitors, Cis-TERN runs a Monte Carlo simulation for up to 10 steps starting with 100 walkers on the scRNA-seq cell of interest. At each step, every walker moves according to row-normalized (row-stochastic) edge weights in *S*^*S*^; rows with all zeros are treated as sinks (walkers remain in that cell). To determine a principled stopping point, Cis-TERN employs an entropy-based termination rule. At each step, it computes the entropy of the walker distribution over cells and applies an elbow criterion: the optimal step is chosen as the one with the largest perpendicular distance to the chord connecting step 1 and the first step at which entropy reaches zero.

### 4.4 Datasets used, data processing, and method run details

#### Data sources

All hematopoiesis (healthy and MPAL) scATAC-seq data was downloaded from GSE139369 as fragment files. scRNA-seq data was obtained from the study’s public GitHub repository as two separate SummarizedExperiment objects: one containing healthy samples, and one for the combined healthy and MPAL dataset. Both objects included curated cell type annotations for all cells in the healthy and MPAL compartments, as provided by the original authors.

#### Preprocessing

The scRNA-seq data was depth-normalized to 10,000 counts per cell to generate the input cell-by-gene expression matrix for Cis-TERN. The scATAC-seq fragment files were processed with ArchR v1.0.2 using default parameters using the functions ‘addArchRGenome(“hg19”)’, ‘createArrowFiles’, ‘addDoubletScores’, ‘ArchRProject’, ‘filterDoublets’, ‘addIterativeLSI’, ‘addHarmony’, ‘addClusers’, ‘addUMAP’, ‘getMarkerFeatures’, ‘addImputeWeights’, ‘addGroupCoverages’, ‘addReproduciblePeakSet’, ‘addPeakMattrix’ in order. ArchR’s binarized cell-by-peak matrix was used as the input cell-by-peak matrix for Cis-TERN. ArchR’s ‘GeneScoreMatrix’ which is a proximal and gene body accessibility-based cell-by-gene matrix that is by default depth-normalized to 10,000 counts per cell, was used as the input cell-by-gene gene score matrix for Cis-TERN. The gene-by-peak peak-gene correlation input matrix was generated by ArchR’s ‘addPeak2GeneLinks’ using the gene integration matrix. The integrated expression matrix was generated by pairing scATAC-seq data with scRNA-seq using ArchR’s ‘addGeneIntegrationMatrix’ function. For the ‘seRNA argument’, we supplied a SummarizedExperiment containing the scRNA-seq data, first subsetting it to include only cells from those samples that were also present in the scATAC-seq dataset, so that integration was performed exclusively between matched samples.

#### Method run details

We ran Cis-TERN on the data in the following combinations: all healthy ATAC with all healthy RNA, one healthy ATAC at a time with all healthy RNA, and one MPAL ATAC and matching RNA with all healthy ATAC and RNA. In each combination, the data preprocessing is run on the given set of data and only genes in common across the two gene-by-cell matrices are kept. In all cases, Cis-TERNs random walk used a damping factor α = 0.85 and a power-iteration convergence tolerance ε = 1×10^−6^. To run Monocle 3, we used the downloaded healthy scRNA-seq SummarizedExperiment and scATAC-seq gene score summarized experiment (generated with ArchR “getMatrixFromProject” with option useMatrix set to “GeneScoreMatrix”). These were converted to Seurat objects using Seurat v5.0.3 function ‘CreateSeuratObject’ (with ‘counts’ set to gene score counts or expression and ‘meta.data’ set to gene score matrix or scRNA-seq SummarizedExperiment column data respectively), log-normalized, and subjected to standard processing with default parameters (highly variable gene selection, scaling and principal component analysis with nPC = 30) [19]. We integrated the scRNA-seq and gene score datasets in gene space using Seurat’s canonical correlation analysis-based integration workflow (‘SelectIntegrationFeatures’, ‘FindIntegrationAnchors’ and ‘IntegrateData’), followed by scaling, PCA and UMAP on the integrated assay. The integrated Seurat object was converted to a Monocle 3 ‘cell_data_set.’ Trajectory learning and pseudotime estimation were performed in Monocle 3 using ‘cluster_cells’, ‘learn_graph’, and ‘order_cells’, with hematopoietic stem cells (annotation ‘01_HSC’ in the ‘BioClassification’ metadata) specified as the root population. The resulting pseudotime values were computed for all cells in the integrated project. For downstream analyses we retained only pseudotime assigned to the scRNA-seq cells.

### 4.5 Code availability

Cis-TERN is an open-source method available at https://github.com/pawelab/Cis-TERN which includes a jupyter notebook with a sample run for pseudotime scoring and reverse tracing functionality.

## Notes

### Competing Interest Statement

The authors have declared no competing interest.

### Funding

- American Cancer Society

## Bibliography

[1] S. L. Klemm, Z. Shipony, and W. J. Greenleaf, “Chromatin accessibility and the regulatory epigenome,” Nat. Rev. Genet., vol. 20, no. 4, pp. 207–220, Apr. 2019.

[2] M. R. Corces et al., “Lineage-specific and single-cell chromatin accessibility charts human hematopoiesis and leukemia evolution,” Nat. Genet., vol. 48, no. 10, pp. 1193–1203, Oct. 2016.

[3] D. Lara-Astiaso et al., “Immunogenetics. Chromatin state dynamics during blood formation,” Science, vol. 345, no. 6199, pp. 943–949, Aug. 2014.

[4] J. D. Buenrostro, P. G. Giresi, L. C. Zaba, H. Y. Chang, and W. J. Greenleaf, “Transposition of native chromatin for fast and sensitive epigenomic profiling of open chromatin, DNA-binding proteins and nucleosome position,” Nat. Methods, vol. 10, no. 12, pp. 1213–1218, Dec. 2013.

[5] M. R. Corces et al., “The chromatin accessibility landscape of primary human cancers,” Science, vol. 362, no. 6413, p. eaav1898, Oct. 2018.

[6] J. D. Buenrostro et al., “Single-cell chromatin accessibility reveals principles of regulatory variation,” Nature, vol. 523, no. 7561, pp. 486–490, July 2015.

[7] J. M. Granja et al., “ArchR is a scalable software package for integrative single-cell chromatin accessibility analysis,” Nat. Genet., vol. 53, no. 3, pp. 403–411, Mar. 2021.

[8] S. Ma et al., “Chromatin potential identified by shared single-cell profiling of RNA and chromatin,” Cell, vol. 183, no. 4, pp. 1103–1116.e20, Nov. 2020.

[9] C. Trapnell et al., “The dynamics and regulators of cell fate decisions are revealed by pseudotemporal ordering of single cells,” Nat. Biotechnol., vol. 32, no. 4, pp. 381–386, Apr. 2014.

[10] L. Heumos et al., “Best practices for single-cell analysis across modalities,” Nat. Rev. Genet., vol. 24, no. 8, pp. 550–572, Aug. 2023.

[11] J. Cao et al., “The single-cell transcriptional landscape of mammalian organogenesis,” Nature, vol. 566, no. 7745, pp. 496–502, Feb. 2019.

[12] Z. Zhang, C. Yang, and X. Zhang, “scDART: integrating unmatched scRNA-seq and scATAC-seq data and learning cross-modality relationship simultaneously,” Genome Biol., vol. 23, no. 1, p. 139, June 2022.

[13] G. Li et al., “Sceptic: pseudotime analysis for time-series single-cell sequencing and imaging data,” Genome Biol., vol. 26, no. 1, p. 209, July 2025.

[14] J. M. Granja et al., “Single-cell multiomic analysis identifies regulatory programs in mixed-phenotype acute leukemia,” Nat. Biotechnol., vol. 37, no. 12, pp. 1458–1465, Dec. 2019.

[15] T. J. Nowakowski et al., “Spatiotemporal gene expression trajectories reveal developmental hierarchies of the human cortex,” Science, vol. 358, no. 6368, pp. 1318–1323, Dec. 2017.

[16] A. E. Trevino et al., “Chromatin and gene-regulatory dynamics of the developing human cerebral cortex at single-cell resolution,” Cell, vol. 184, no. 19, pp. 5053–5069.e23, Sept. 2021.

[17] E. Becht et al., “Dimensionality reduction for visualizing single-cell data using UMAP,” Nat. Biotechnol., vol. 37, no. 1, pp. 38–44, Dec. 2018.

[18] H. Chen et al., “Assessment of computational methods for the analysis of single-cell ATAC-seq data,” Genome Biol., vol. 20, no. 1, p. 241, Nov. 2019.

[19] Y. Hao et al., “Dictionary learning for integrative, multimodal and scalable single-cell analysis,” Nat. Biotechnol., vol. 42, no. 2, pp. 293–304, Feb. 2024.

