## Appendix for "Cis-TERN: a graph-based method to integrate single-cell chromatin accessibility with gene expression data to infer the pseudotime ordering of cells"

The appendix contains the following additional figures:

Figure 1: Overview of reverse tracing with Cis-TERN
Figure 2: HSCs have an inflated number of incoming non-zero edges
Figure 3: Cis-TERN scores for all cell types in healthy hematopoiesis
Figure 4: Cis-TERN orders cell lineages in brain development
Figure 5: Comparison of Cis-TERN scores to UMAP embedding
Figure 6: Robustness of Cis-TERN to smaller quantities of scATAC-seq cells
Figure 7: Cis-TERN runtime scaling
Figure 8: Evaluation of precedence scores and cosine similarity
Figure 9: Cancer data does not reorder healthy cells
Figure 10: Reverse tracers make spurious visits
Figure 11: Distribution of trace lengths before termination
Figure 12: Reverse tracing of cancer cells converges rapidly

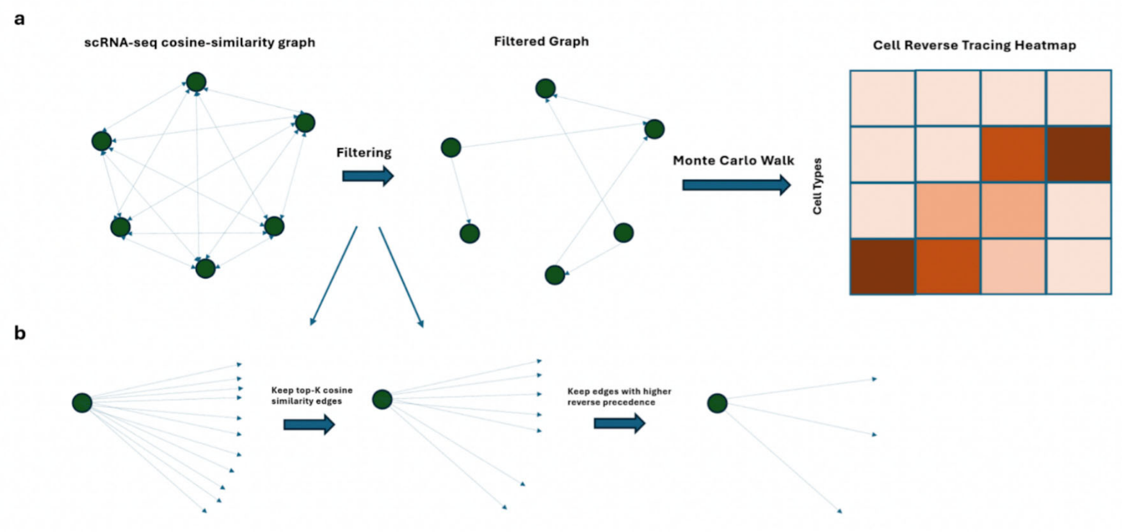


Appendix Figure 1. **Overview of reverse tracing with Cis-TERN. a)** For reverse tracing, cis-TERN builds a pairwise cosine similarity matrix of scRNA-seq cells from which it filters edges out. It then runs a Monte Carlo simulation for up to 10 steps starting with 100 walkers on the scRNA-seq cell of interest. At each step, every walker moves according to row-normalized (row-stochastic) edge weights. **b)** Cis-TERN filters edges per cell by first keeping only the top 10% of outgoing edge weights and then removing edges to any cells that have a greater reverse Cis-TERN score.

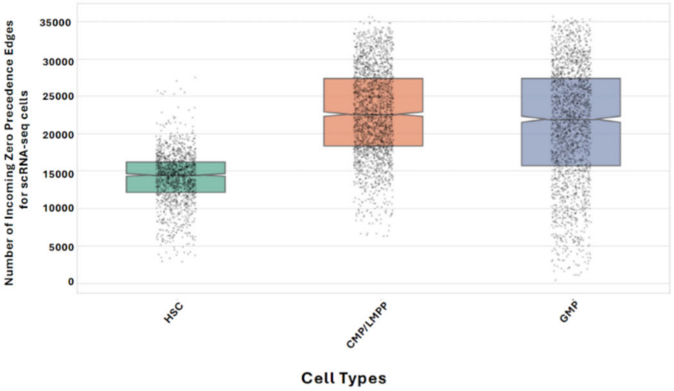


Appendix Figure 2. **HSCs have an inflated number of incoming non-zero edges.** Notched boxplots of the number of incoming zero-weighted precedence edges for scRNA-seq cells in three early progenitor cell types.

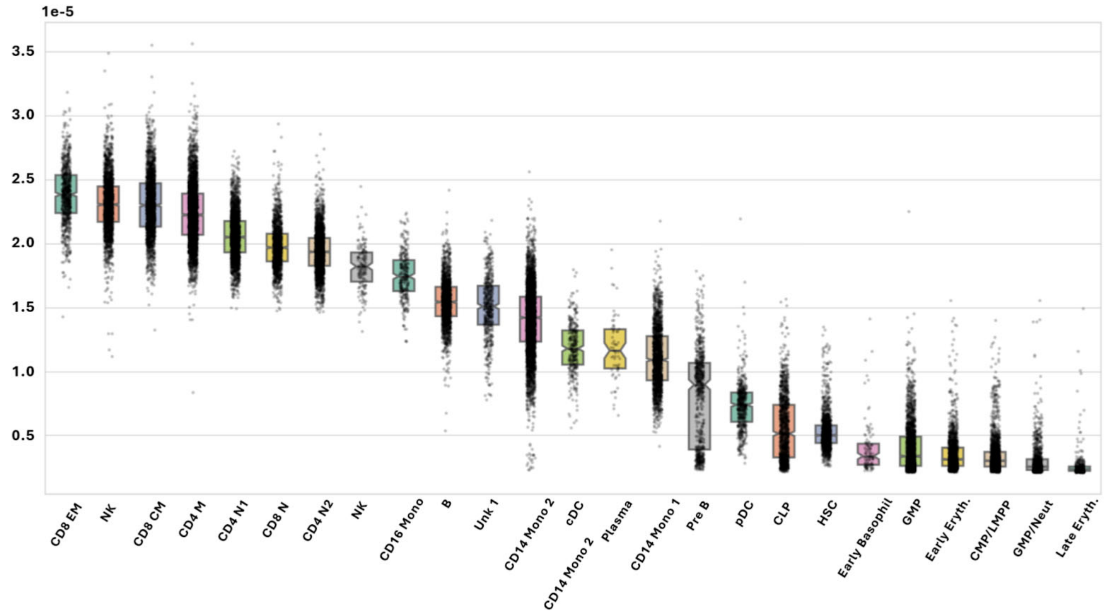


Appendix Figure 3. **Cis-TERN scores for all cell types in healthy hematopoiesis.** Notched boxplots of Cis-TERN scores for all cell types in healthy hematopoietic data ordered by median Cis-TERN score.

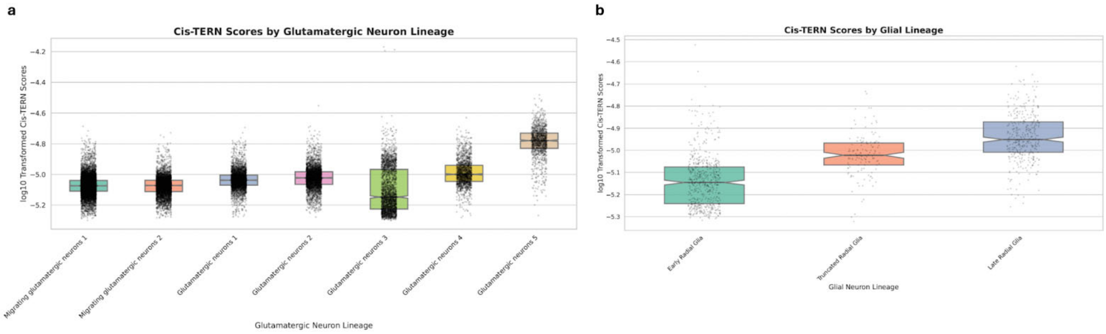


Appendix Figure 4. **Cis-TERN orders cell lineages in brain development.** Notched boxplots of Cis-TERNs scores for cells in **a)** the glutamatergic neuron lineage **b)** the radial glia lineage. For this analysis, scATAC-seq data was downloaded as fragment files from GSE162170 and scRNA-seq data was obtained from the same study's public GitHub repository as a SummarizedExperiment [16]. The SummarizedExperiment included curated cell type annotations. Overall the dataset contained 57,868 scRNA-seq cells and 31,304 scATAC-seq cells from four fetal cortex samples (PCW16, 20, 21, 24) and the scRNA-seq cells were annotated into 23 cell types. The preprocessing steps and Cis-TERN run details were the same as described in Methods section 4.4.

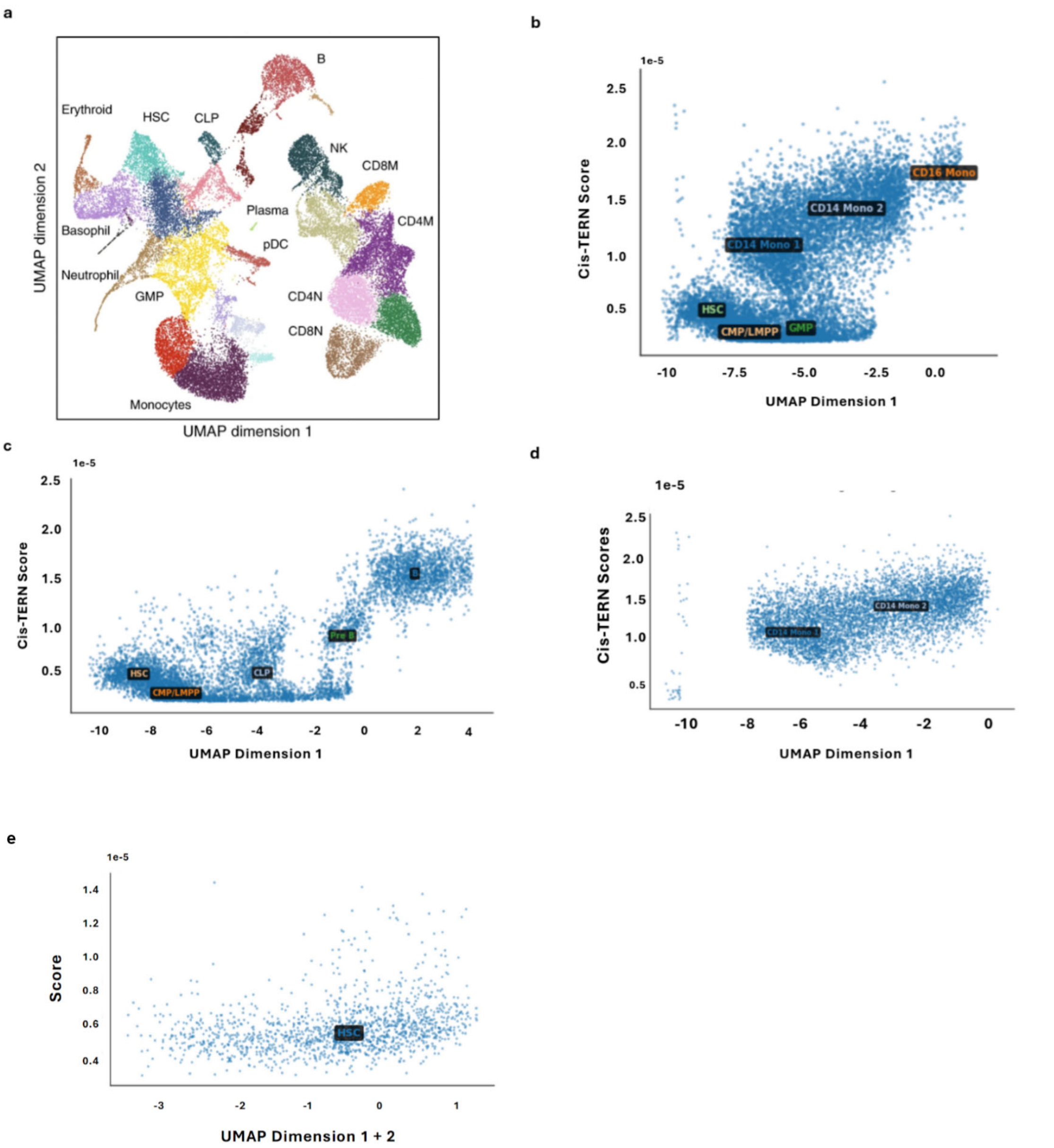


Appendix Figure 5. **Comparison of Cis-TERN scores to UMAP embedding. a)** UMAP embedding of all healthy hematopoietic scRNA-seq cells [14]. Cis-TERN score vs UMAP Dimension 1 for **b)** the monocyte cell lineage **c)** the B-cell lineage and **d)** CD14 Mono 1 and CD14 Mono 2 cells. **e)** Cis-TERN score vs UMAP Dimension 1 plus Dimension 2 for HSCs. Cell type labels appear at the centroid of all cells assigned to that cell type.

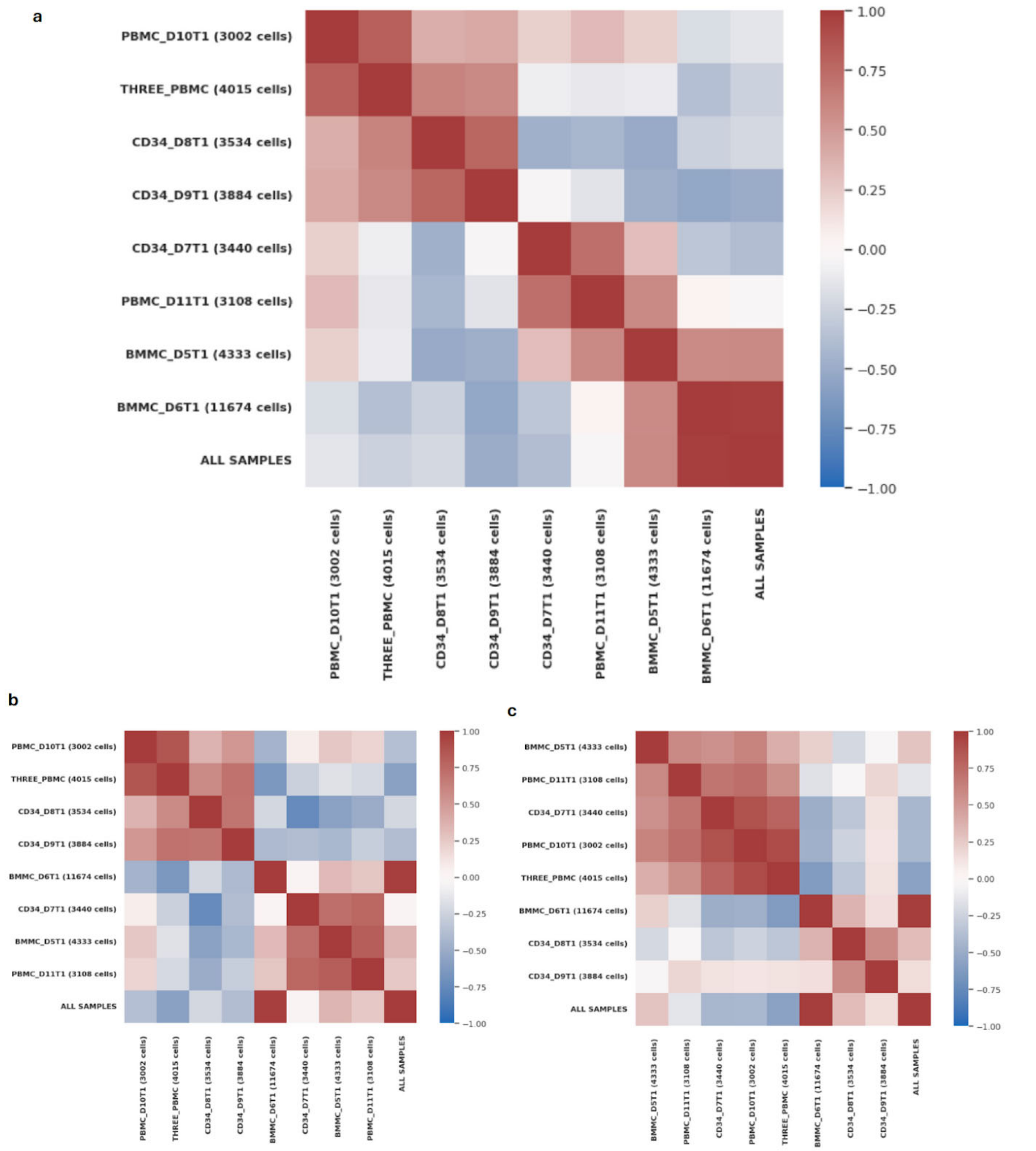


Appendix Figure 6. **Robustness of Cis-TERN to smaller quantities of scATAC-seq cells.** Heatmaps of the Pearson correlations of Cis-TERN scores for **a)** all scRNA-seq cells **b)** cells in the monocyte lineage and **c)** cells in the B-cell lineage when using only the given sample's scATAC-seq cells. While samples with more than 4,000 cells remained highly correlated, performance degraded below that point.

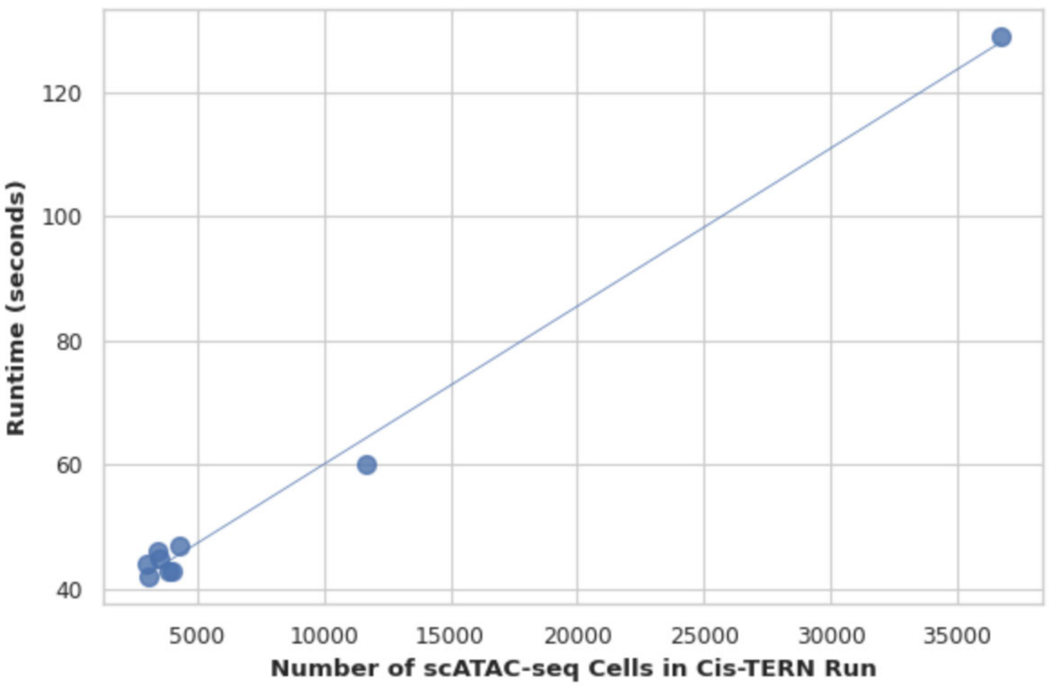


Appendix Figure 7. **Cis-TERN runtime scaling.** Runtime of Cis-TERN in seconds for varying numbers of scATAC-seq cells. scRNA-seq cells are kept constant to 35,882.

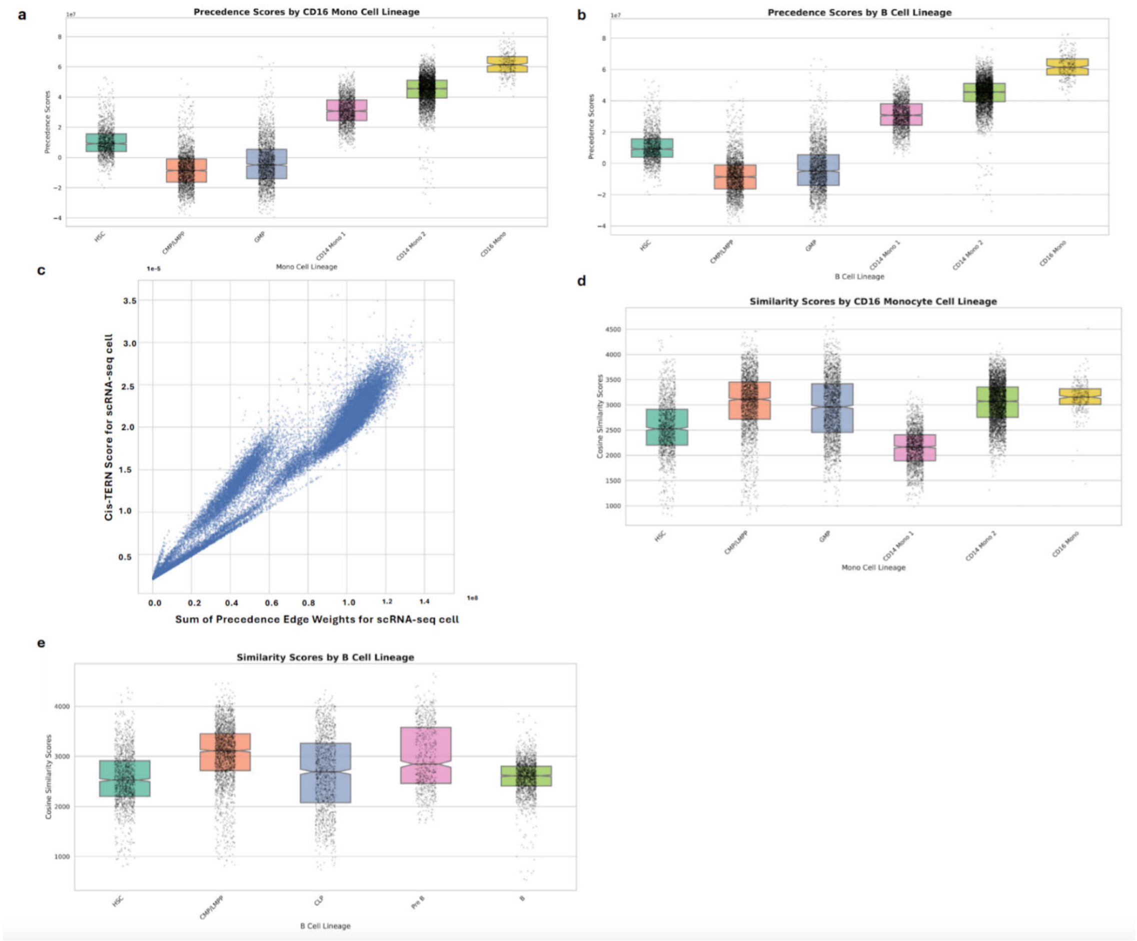


Appendix Figure 8. **Evaluation of precedence scores and cosine similarity.** Notched boxplots of the sum of incoming precedence scores for scRNA-seq cells in **a)** the monocyte lineage and **b)** the B-cell lineage. **c)** Comparison of summed precedence scores to cis-TERN scores. Notched boxplots of Cis-TERN scores when replacing precedence scores with cosine similarity in the graph for **a)** the monocyte lineage and **b)** the B-cell lineage results in a loss of ordering of cell types in each lineage.

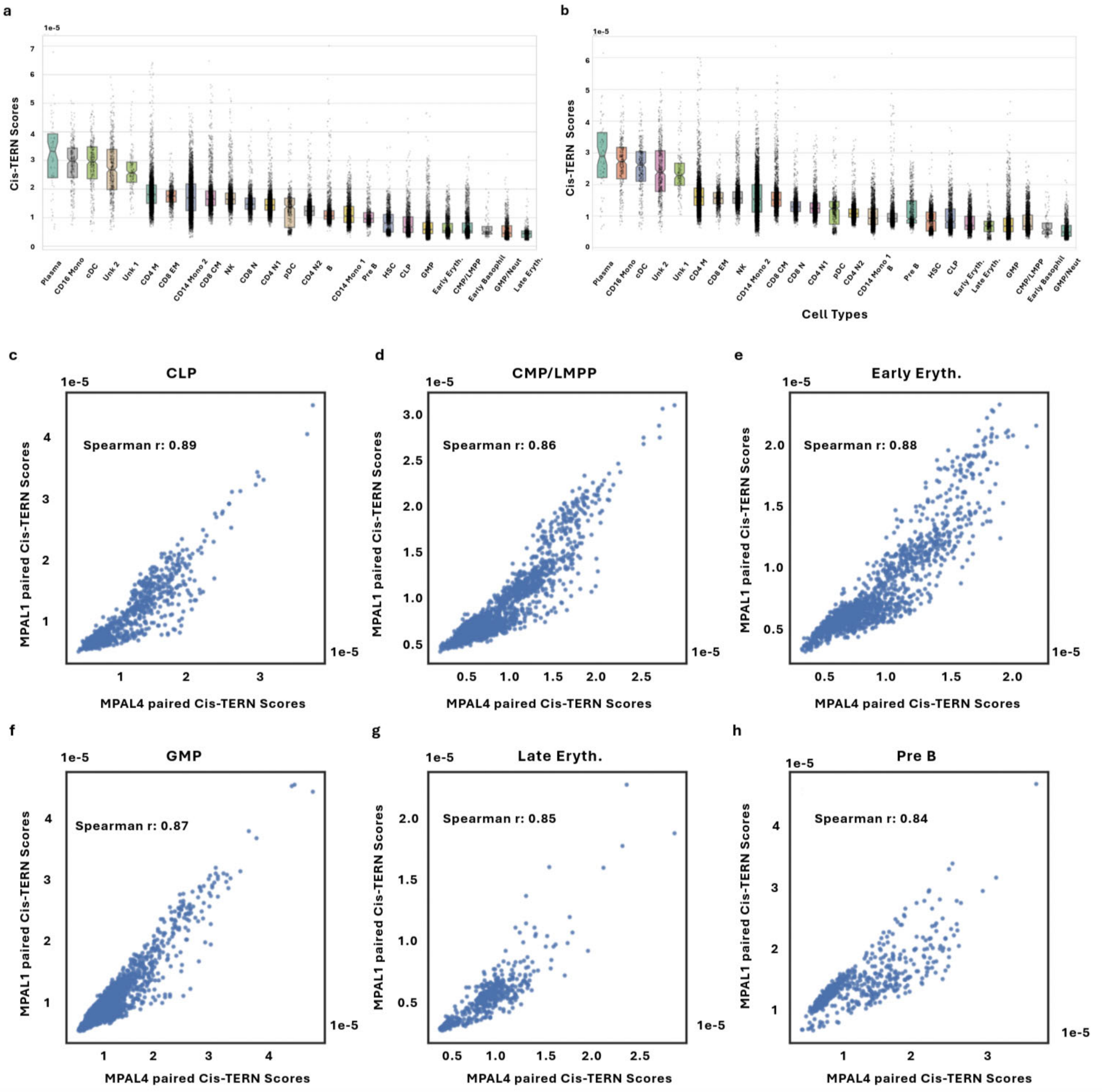


Appendix Figure 9. **Cancer data does not reorder healthy cells.** Cis-TERN scores for healthy cell types when paired with **a)** MPAL1 (T-myeloid MPAL) **b)** MPAL4 (B-myeloid MPAL). Per cell type, cis-TERN scores were highly correlated between MPAL1 and MPAL4 with all Spearman's $\rho > 0.9$ except for **c)** CLP cells, **d)** CMP/LMPP cells, **e)** early erythrocytes, **f)** GMP cells, **g)** late erythrocytes, and **h)** pre-B cells, all of which had Spearman's $\rho > 0.8$.

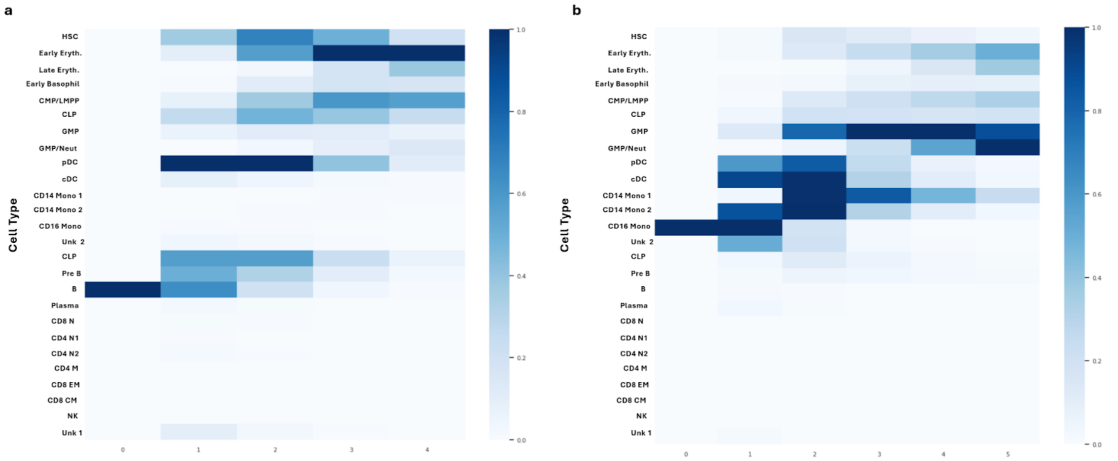


Appendix Figure 10. **Reverse traces make spurious visits.** The min-max scaled fraction of traces that visited each cell type when traces started at **a)** B cells and **b)** CD16 monocytes. The traces make additional spurious visits to cell types outside of the known lineages.

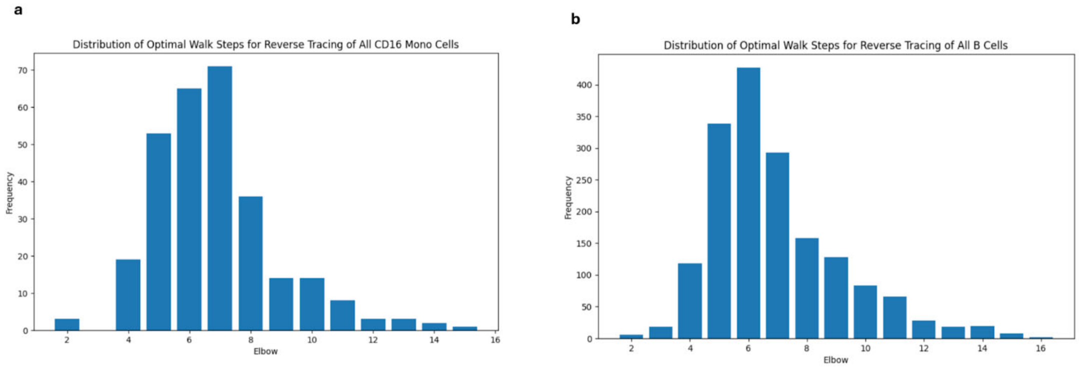


Appendix Figure 11. **Distribution of trace lengths before termination.** The distribution of optimal walk steps for reverse tracing of **a)** all CD16 Mono cells and **b)** all B cells based on elbow method termination.

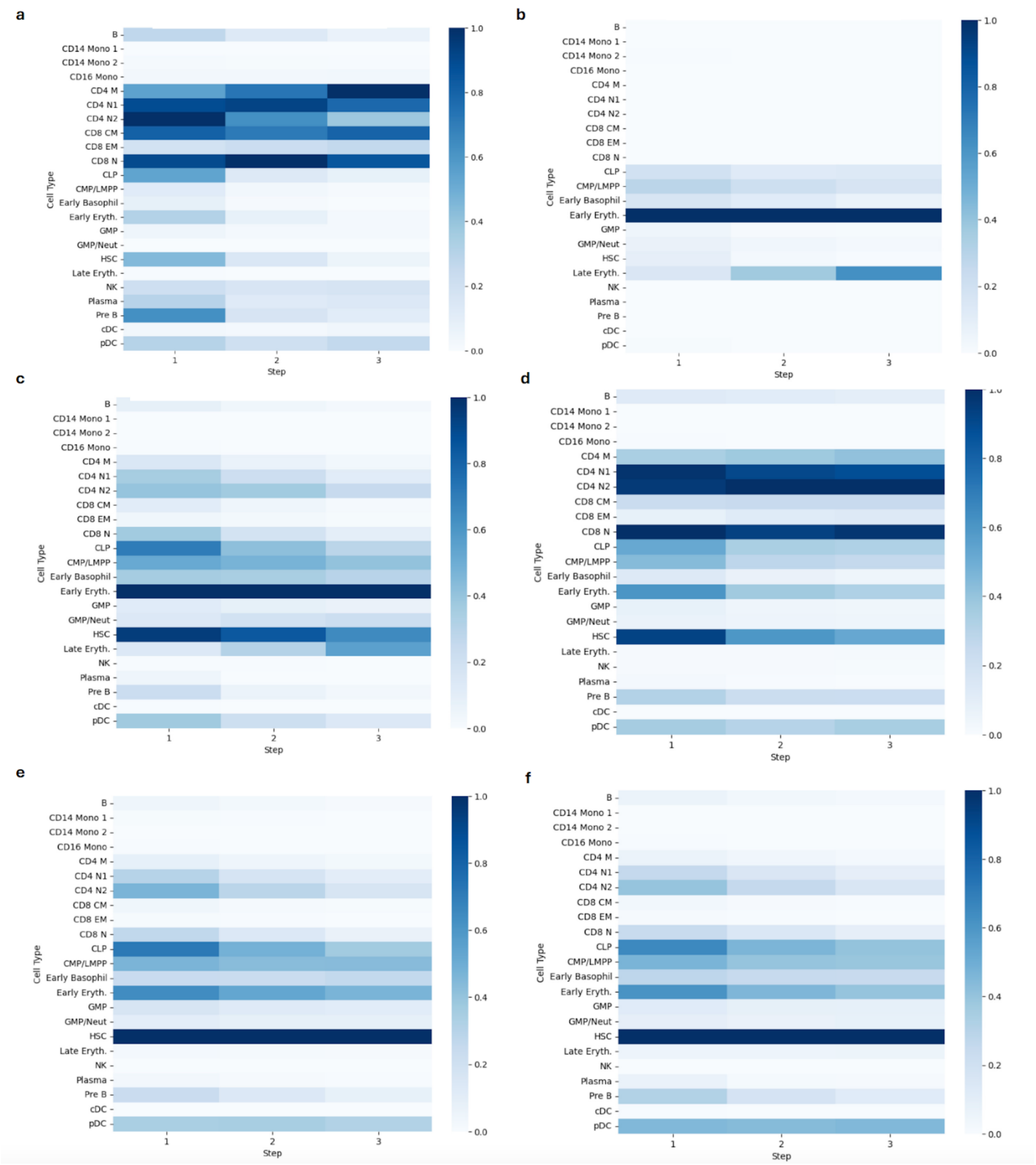


Appendix Figure 12: **Reverse tracing of cancer cells converges rapidly.** Cis-TERN reverse tracing results for starting with all cancer cells at step 0 for **a)** MPAL1, **b)** MPAL2, **c)** MPAL3, **d)** MPAL4, **e)** MPAL5, and **f)** MPAL5R converge after three steps.
